# Claw4Science: A Dataset and Platform for the OpenClaw Scientific Agent Ecosystem

**DOI:** 10.64898/2026.03.30.715118

**Authors:** Mingyang Xu, Junhao Chen, Zaixi Zhang

## Abstract

Large language models have enabled a new class of scientific software in the form of AI agents that can execute research workflows across bioinformatics, drug discovery, and related domains. Among these systems, OpenClaw introduced a skill-based design that allows workflows to be expressed as structured Markdown files, lowering the barrier to contribution and enabling rapid ecosystem growth. However, this growth has led to fragmentation. Projects are distributed across independent repositories, skills vary widely in quality, naming is inconsistent, and there is no unified way to discover or compare available tools. In this work, we construct the first curated dataset of the OpenClaw scientific ecosystem. The dataset includes 91 projects organized by functional role and 2,230 skills spanning 34 scientific categories. Based on this dataset, we perform a systematic analysis of the structure, distribution, and emerging patterns of scientific agent development. To make this ecosystem accessible in practice, we further build Claw4Science, a public platform at https://claw4science.org, which is built on top of our dataset. The platform organizes projects and aggregates distributed skill repositories into a unified interface, with a focus on bioinformatics and scientific workflows, providing a practical entry point for navigating the ecosystem. Our results show that the OpenClaw ecosystem reflects a shift from isolated systems to a more modular and shareable model of scientific computation. At the same time, challenges in evaluation, reproducibility, and governance remain open. We argue that our dataset provides a foundation for future benchmark development and standardized infrastructure for scientific AI agents.

## 1. Introduction

The idea of automating scientific research has a long history, but for most of that time there was a clear gap between what people hoped for and what systems could actually achieve. Early closed-loop discovery systems showed that machines could propose and test hypotheses without continuous human intervention. However, these systems worked only in narrow and carefully designed settings. Systems such as AutoBA, CellVoyager, BioDiscoveryAgent, and the Virtual Lab show that a general-purpose language model, when combined with appropriate tools and scaffolding, can perform useful tasks across RNA-seq analysis, genetic perturbation design, spatial transcriptomics, and protein engineering [35, 45, 76]. Recent surveys report steady progress in applying these systems to chemistry, biology, and materials science [44, 54]. At the same time, the idea of AI as a scientific collaborator has gained attention in major venues [53, 60].

These early systems were important, but they shared a key limitation. Each system was largely self-contained. Capabilities were implemented in source code. Extending a system required modifying that code. Sharing a new workflow usually meant releasing a new repository or publishing a paper. As a result, adapting existing systems was diffcult. A researcher who wanted to adapt CellVoyager’s single-cell analysis logic for a different organism, or combine AutoBA’s omics pipeline with a custom visualization step, had to deal with software engineering challenges rather than scientific ones. There was no simple way to share and reuse workflows across systems. The community lacked a lightweight mechanism to accumulate and redistribute scientific knowledge. As a result, these agents remained isolated and did not form a broader ecosystem[71].

OpenClaw addresses this problem at the architectural level[7]. It introduces the concept of a skill as a core extension unit. A skill is a structured Markdown file that contains natural language instructions and optional executable code blocks[2]. Any compatible agent runtime can read and execute it. Writing a skill does not require modifying the underlying agent code or a software engineering background. A computational biologist can write a skill in the same way they would write a protocol. This design makes it easier to build, extend, and share agent capabilities. The same skill can run across multiple OpenClaw-compatible systems without modification. OpenClaw also integrates gateway management, persistent memory, and multi-channel messaging in a single system, lowering the barrier to developing scientific agents. As a result, the ecosystem expanded rapidly. Independent teams released domain-specific extensions and skill libraries across bioinformatics, clinical research, and general scientific workflows[15, 59, 69, 73]. As of March 2026, the ecosystem includes more than 91 projects with over 50 GitHub stars each, accumulating over 450,000 stars, and 2,230 curated science skills across 34 categories[74]. This rapid expansion also makes it diffcult to systematically track and compare the ecosystem, motivating the need for a structured dataset.

This rapid growth is a clear success, but it also introduces new problems. Projects appear faster than researchers can track, and there is no unified directory or navigation resource. Most projects are distributed across independent repositories, making it diffcult to identify suitable tools. Name collisions are a concrete issue. Multiple independent projects share identical names such as “ScienceClaw” or “PaperClaw,” which leads to confusion[33, 34, 36, 46, 58, 59, 69, 75]. The skill layer also lacks a standardized evaluation framework. The 2,230 skills vary widely in quality, but there is no principled way to compare them[4, 14]. In addition, there is no clear method to evaluate the scientific capability of agents. Claims about self-evolving systems remain diffcult to verify without agreed benchmarks. This paper addresses these challenges and provides a structured view of the ecosystem. Our contributions are as follows. Together, these issues make it diffcult to form a coherent view of the ecosystem.

This work provides a structured view of the OpenClaw scientific ecosystem through a dataset, an analysis, and a platform. We construct a curated dataset that includes 91 projects organized by functional role and over 2,200 skills spanning 34 scientific categories. Based on this dataset, we analyze the structure of the ecosystem, including its functional taxonomy, skill distribution, and the emerging role of skills as a unit of scientific contribution. We also examine early evaluation efforts such as the Claw4S Conference [3], and identify key challenges, including naming collisions, skill quality variation, and the lack of standardized benchmarks. To make this ecosystem accessible in practice, we build Claw4Science (https://claw4science.org), a platform that organizes projects and aggregates distributed skill repositories into a unified interface for scientific workflows. Finally, we highlight open directions, including evaluation frameworks, governance, and edge-deployed scientific agents.

## 2. Scientific AI Agents Before OpenClaw

The projects that preceded the OpenClaw ecosystem were often technically strong. AutoBA showed that a language model agent could plan and execute multi-omic analyses across RNA-seq, single-cell sequencing, and whole-genome workflows with minimal user input, while recent systems such as STELLA further explore multimodal and tool-augmented scientific agents [35, 45, 49, 76]. CellVoyager explored single-cell datasets and generated hypotheses beyond standard pipelines[35]. BioDiscoveryAgent treated genetic perturbation design as a closed-loop reasoning problem and improved over Bayesian optimization baselines[64]. In molecular science, ChemCrow applied language model agents to tasks in synthesis, drug discovery, and materials design[56]. The Virtual Lab used multiple agents to design nanobodies against SARS-CoV-2 variants and validated the results experimentally[68]. CRISPR-GPT automated guide RNA design and gene-editing planning [63]. ChemGraph coordinated computational chemistry and materials workflows through an agent-based system [61]. Together, these systems demonstrate the growing capability of language model agents in scientific workflows across multiple domains.

These systems addressed real scientific bottlenecks. They reduced the expertise needed to run complex pipelines and helped accelerate hypothesis generation. In some cases, they produced results that were published in high-impact venues. However, none of them led to sustained community growth. The key issue was not what these systems could do, but how they could be extended and shared. Each of these agents was a closed system. Capabilities were implemented in source code. Tool integrations, prompt templates, workflow logic, and domain knowledge were all embedded in the repository. Adding a new step or adapting a pipeline required modifying the code. Combining capabilities across systems required the same effort. For most domain scientists, this was a major barrier. They needed to understand the codebase, modify it safely, and maintain their changes over time. As a result, systems evolved slowly and were mainly developed by their original authors or a small group of technical contributors. Adoption increased through citations and GitHub stars, but the community could not easily build on these systems or reuse their workflows. This also makes it diffcult to systematically compare and study these systems across domains.

There was no shared substrate across these systems. AutoBA’s omics logic could not be transferred to CellVoyager’s single-cell framework without substantial rewriting. A CRISPR-GPT workflow could not be combined with an MDCrow simulation pipeline without custom integration [38]. Each agent used its own format and runtime. As a result, the community produced many capable but isolated tools. Each system solved a specific problem well, but there was no mechanism for these solutions to accumulate into a larger ecosystem. This is not a criticism of these systems. They addressed diffcult problems, and their design choices were reasonable at the time. The issue is structural. The pre-OpenClaw landscape lacked a shared layer for agent capabilities. There was no simple and portable way for a scientist to encode a workflow and share it across systems. There was also no common runtime that non-engineers could extend without modifying source code. As a result, the knowledge embodied in these agents remained isolated, and the ecosystem was diffcult to study or compare in a systematic way. This structural gap is what the OpenClaw skill system addresses. It introduces a portable and shareable unit of workflow representation, which enables the emergence of a broader ecosystem. The next section examines this in detail. We provide a visual summary of this transition in Figure 1.

**Figure 1.**
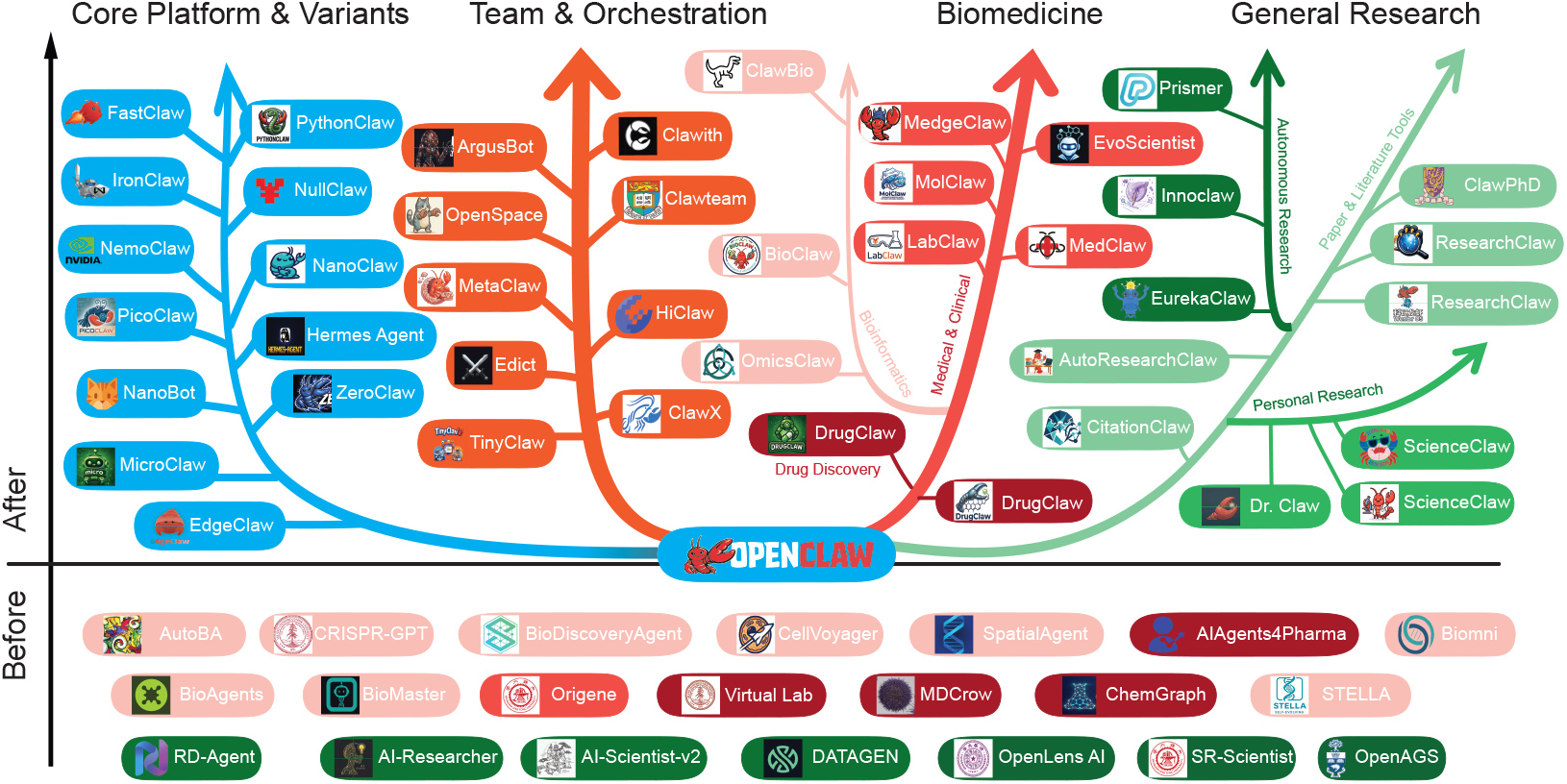
Overview of the OpenClaw scientific ecosystem. The figure organizes projects into major branches, including core platform variants, team and orchestration systems, biomedicine-related agents, and general research agents. It also highlights the shift from pre-OpenClaw systems to a broader post-OpenClaw ecosystem.

## 3. Dataset-Driven Ecosystem Analysis

### 3.1. Scope and Classification Logic

Our analysis is based on a curated dataset of OpenClaw-related projects and skills. Figure 1 and Figure 2 provide two complementary views of the post-OpenClaw ecosystem. The first shows a functional taxonomy, while the second shows the structural relationships between projects. The functional taxonomy groups projects by their main purpose. We assign each project to the role it primarily serves in practice, including core runtimes, orchestration systems, domain-specific scientific agents, and skill libraries or research workspaces. Some projects support multiple functions, but we classify them by their central design goal.

**Figure 2.**
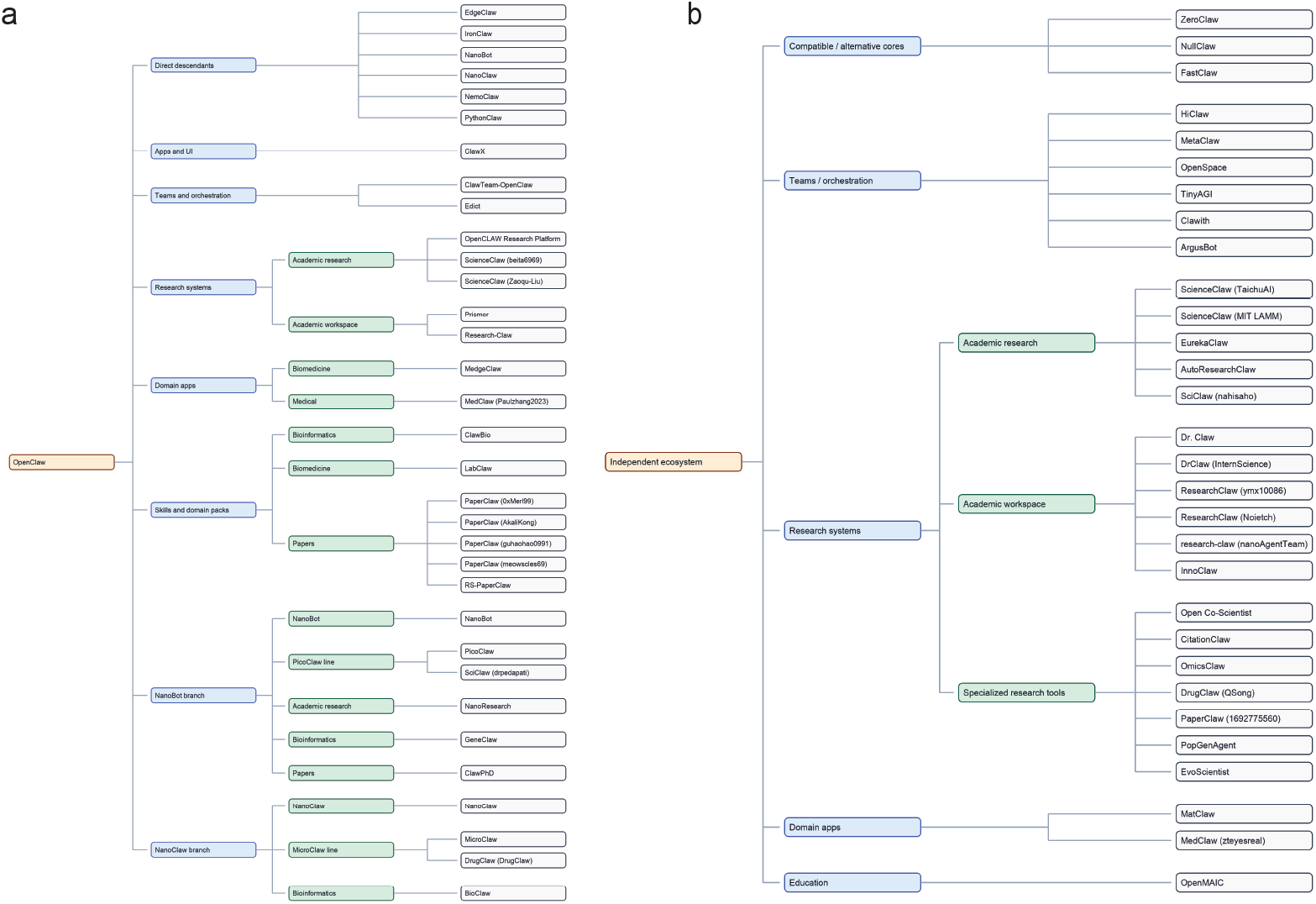
Structural map of the post-OpenClaw ecosystem. The figure distinguishes direct OpenClawderived branches from adjacent independent systems, and organizes projects into major layers such as core platform variants, orchestration systems, research systems, domain applications, and skill libraries. Unlike Figure 1, this map is intended to provide a more systematic view of ecosystem structure rather than a purely conceptual overview.

**Figure 3.**
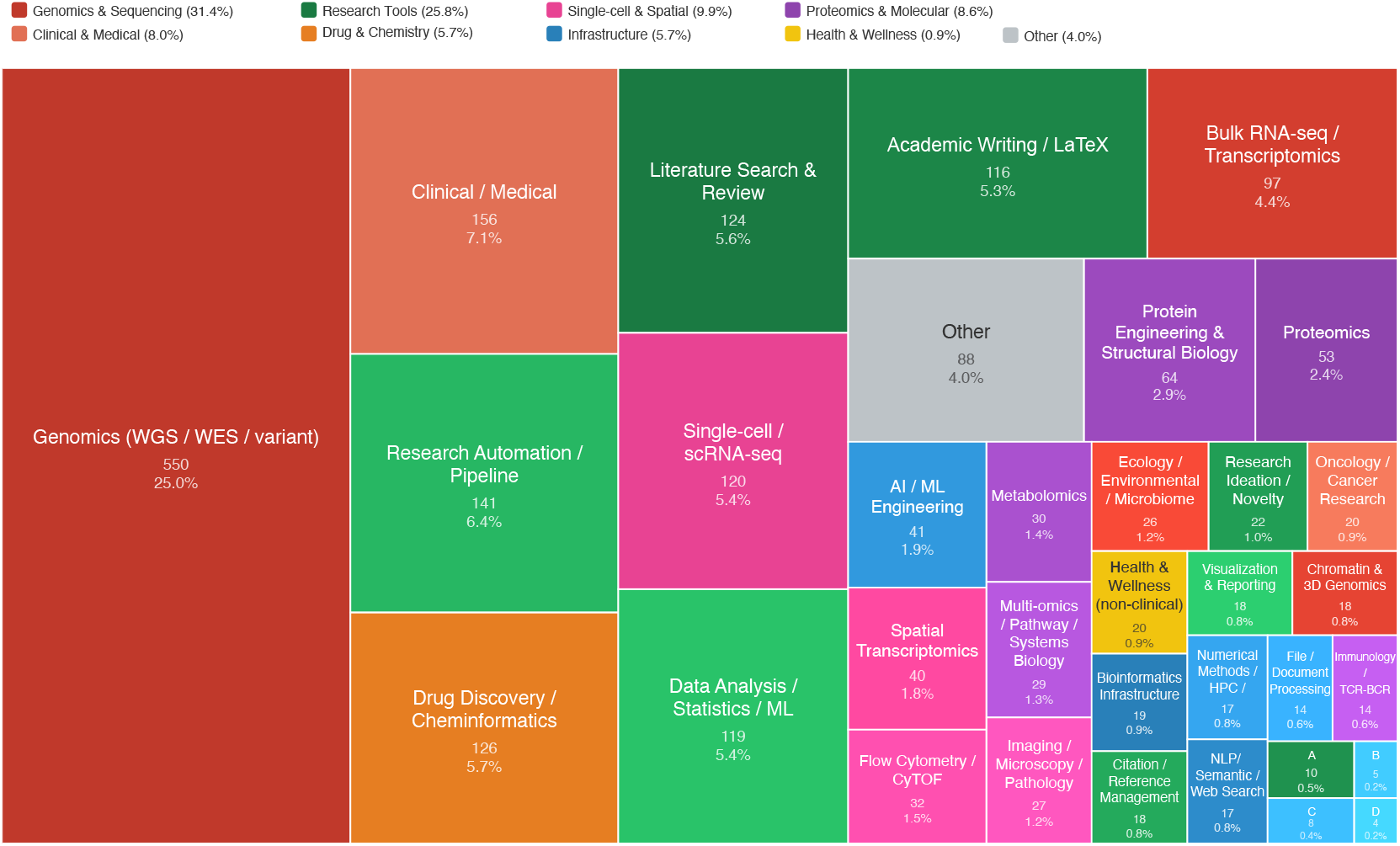
Distribution of 2,230 skills across 34 scientific categories. Area represents the number of skills in each category. Due to space constraints, several minor categories are abbreviated as (A) research review and peer review, (B) physics, materials, and earth sciences, (C) general and developer tools, and (D) finance and economics.

The structural view focuses on how each project relates to OpenClaw. Some projects are direct descendants. Others are lightweight variants or specialized extensions. Some are independent systems that remain compatible with or influenced by OpenClaw. We keep these two views separate. A project may belong to one functional category but occupy a different structural position. This distinction is important because similar names do not imply a shared technical base. Our classification is pragmatic. We group projects by primary function and note structural relationships when they are clear enough to support a reliable judgment.

### 3.2. Ecosystem Dataset and Data Collection

We construct a curated dataset of OpenClaw-related projects and skills through a multi-source discovery and manual review process. Candidate projects were identified via GitHub keyword search, social media monitoring, community recommendations, and cross-references from existing repositories. Each project was manually reviewed to ensure relevance to scientific workflows and public availability. In total, we collected 62 OpenClaw-related projects released after the introduction of the platform. These projects form the basis of the ecosystem analysis in this work. In addition to project-level data, the dataset includes a large-scale view of the skill layer, which serves as the main extension mechanism of the ecosystem. We developed an automated pipeline to scan 12 science-focused skill repositories via the GitHub API. This process identified over 2,200 skills and classified them into 34 domain categories using keyword-based matching, with a 96.0% classification rate. Project metadata, including GitHub stars and activity status, is updated regularly. The dataset is maintained as a living resource at https://claw4science.org.

### 3.3. Two Complementary Views of the Ecosystem

Figure 1 and Figure 2 provide complementary views of the ecosystem based on our dataset. Figure 1 gives a conceptual overview, highlighting the main branches and the overall shape of post-OpenClaw development. Figure 2 provides a more detailed structural map. It distinguishes direct OpenClaw branches from adjacent systems and shows how platform variants, orchestration layers, research systems, and domain applications are organized. Together, the two figures provide both a high-level view and a more precise structural understanding of the ecosystem.

### 3.4. Core Platform and Structural Layers

Based on our dataset, the post-OpenClaw ecosystem does not appear as a flat list of tools, but as a layered structure, as shown in Figure 1 and Figure 2. At the center is the core platform layer. Around it are lightweight runtime variants, orchestration systems, research workspaces, skill libraries, and domain applications. This layered structure helps explain the rapid expansion of the ecosystem. OpenClaw forms the core of this structure. It established key conventions such as messaging-first interaction, multi-channel use, and a skill-based extension model. These choices lower the cost of adapting a general agent to different research settings. In many cases, new systems can reuse the same runtime and extend it with skills, tools, or workflows.

Around this core, lightweight variants emerge. These systems keep the basic OpenClaw model but optimize for different goals. Some focus on simplicity and local use, while others emphasize security or deployment flexibility[8, 20, 23–25, 27, 32]. Together, they show that OpenClaw has become a base that others reimplement, simplify, or specialize. This platform layer enables a second layer of orchestration and interface systems. These projects move beyond single-agent use and support coordination between agents or interaction through graphical interfaces[31, 39, 62]. They treat OpenClaw as infrastructure rather than as a complete application.

A third layer consists of research systems and workspaces. Some are directly built on OpenClaw, while others are structurally independent but play similar roles[12, 13, 16, 19, 24, 32]. This highlights the need for a structural view, since systems that appear similar at the functional level may differ in their technical foundations. A fourth layer includes skill libraries and domain packs. These projects extend existing runtimes with domain knowledge and workflow templates. This is especially important in scientific contexts, where many tasks depend on reliable procedures rather than new architectures[15, 69, 73]. The skill system makes these procedures easier to share and reuse. Finally, domain applications sit on top of these layers. These systems apply the same underlying structure to specific scientific areas such as biomedicine or drug discovery[17, 21, 42]. Many of them build not only on OpenClaw itself, but also on its downstream variants.

Overall, this layered pattern is consistently observed in our dataset. OpenClaw matters not only as a single project, but as a shared substrate. Some systems inherit its structure directly, while others adapt or extend it. Together, these layers turn a single agent framework into a broader scientific ecosystem.

### 3.5. Scientific Application Layers

Based on our dataset, the scientific application layer builds on the platform and orchestration layers shown in Figure 1 and Figure 2. This is where the OpenClaw ecosystem connects most directly to research practice. Instead of only producing runtimes or coordination tools, the ecosystem supports agents and workflows for real scientific tasks.

One group of projects focuses on general research automation. These systems aim to support the full research process, including literature review, planning, coding, analysis, and writing[5, 10, 11, 18, 22, 29, 55, 58]. Some are built directly on OpenClaw, while others are structurally independent but follow similar design ideas. Together, they reflect a broader shift toward agent-based research workflows.

Another group focuses on domain-specific scientific applications. In bioinformatics and omics, many projects support genomics, transcriptomics, and related analyses[30, 37, 40, 66]. These workflows are data-intensive and require strong integration across tools. In drug discovery and molecular research, other projects focus on evidence retrieval, reasoning, and simulation-driven tasks[17, 67]. Although these domains differ in tooling, they adopt a similar agent-based structure. A smaller group covers education and other specialized domains[9]. These projects show that the ecosystem is not limited to a single research area, but can be adapted to different scientific settings.

Overall, this pattern is consistently observed in our dataset. The ecosystem expands from general-purpose agents to a broader set of scientific tools. Many of these tools share common workflows and runtime ideas. The growth of the ecosystem is therefore not only an increase in project count, but also a shift toward domain-specific scientific applications.

### 3.6. Cross-Cutting Observations

Based on our dataset, several patterns emerge across the ecosystem. First, attention is highly concentrated. A small number of core platform projects account for most of the stars, while many science-facing projects remain in the long tail. Visibility and scientific relevance do not always align. Second, there is a clear split between platform development and scientific applications. Core runtimes and lightweight variants receive most attention, while many science-oriented tools are smaller and more specialized. This suggests that OpenClaw mainly serves as a shared base for downstream tools. Third, programming language choice follows function. Core platforms use a mix of TypeScript, Python, Rust, and Go[20, 25, 27, 28]. Science-facing tools are mostly written in Python, which is consistent with scientific computing practice[15, 30]. This reflects different priorities in system design.

Fourth, institutional participation is uneven. Some projects are linked to universities or research labs[9], while many others are developed by small teams or individual contributors. The ecosystem combines formal research efforts with informal open-source work. Fifth, naming is often unreliable. Similar names can refer to different systems with unrelated technical bases. This makes structural analysis necessary to avoid confusion. Finally, the growth of scientific tools is real but uneven. More agents are being built and shared, but most remain small and lightly validated. Overall, the ecosystem is better understood as a growing research tool landscape than as a mature platform. These patterns are consistently observed in our dataset.

## 4. The Skill System: The Engine of the Ecosystem

### 4.1. What Is a Skill?

A skill is the basic unit of the OpenClaw ecosystem[4]. It is a structured Markdown file that defines a reusable analytical workflow. The file includes metadata such as a name, a description, a type, and trigger conditions[14]. It also contains step-by-step instructions written in natural language, with optional code blocks in Python, Bash, R, or other languages. Any compatible agent runtime can read and execute it. Writing a skill does not require modifying the agent code. It also does not require a software engineering background. A researcher can write a skill in the same way they would describe a protocol. The entire workflow is stored in a single text file that can be read, edited, and version-controlled with standard tools.

This format differs from traditional extensibility mechanisms. Plugin systems require contributors to write code against platform-specific APIs [43]. Workflow systems such as Galaxy reduce this burden by providing graphical interfaces, but workflows are tied to a specific platform [70]. Jupyter notebooks combine code and narrative, but they depend on a specific execution environment [50]. In contrast, a skill is not tied to a single system. Any LLM-based runtime that can read Markdown and execute code blocks can use it. The same analysis skill can run on OpenClaw, NanoBot, PicoClaw, and Claude Code without modification[2, 7, 20, 28]. This portability reduces fragmentation across tools and makes it easier to share workflows between systems. Based on our dataset, the skill layer emerges as the primary mechanism for extending functionality across the ecosystem. Recent work on prompt templates shows that many LLM applications follow similar structural patterns [57]. A skill can be seen as a structured and reusable version of this idea, designed for scientific workflows. Unlike ad hoc prompts, skills are documented, versioned, and easier to reuse. They encode the practical knowledge of a researcher in a form that others can directly apply.

### 4.2. The Skill Ecosystem at Scale

Based on our dataset, Figure 4.1 shows the distribution of 2,230 skills across 34 categories from claw4science Skill Hubs [4, 6, 26, 65, 72, 74]. The distribution is highly uneven. A small number of categories dominate. Genomics accounts for the largest share, followed by clinical and medical applications, drug discovery, and research automation. Together, these categories make up a large fraction of the ecosystem. At the same time, there is a long tail of smaller categories. These include metabolomics, imaging, citation management, and visualization. Each category is small on its own, but together they cover a wide range of scientific tasks.

**Figure 4.**
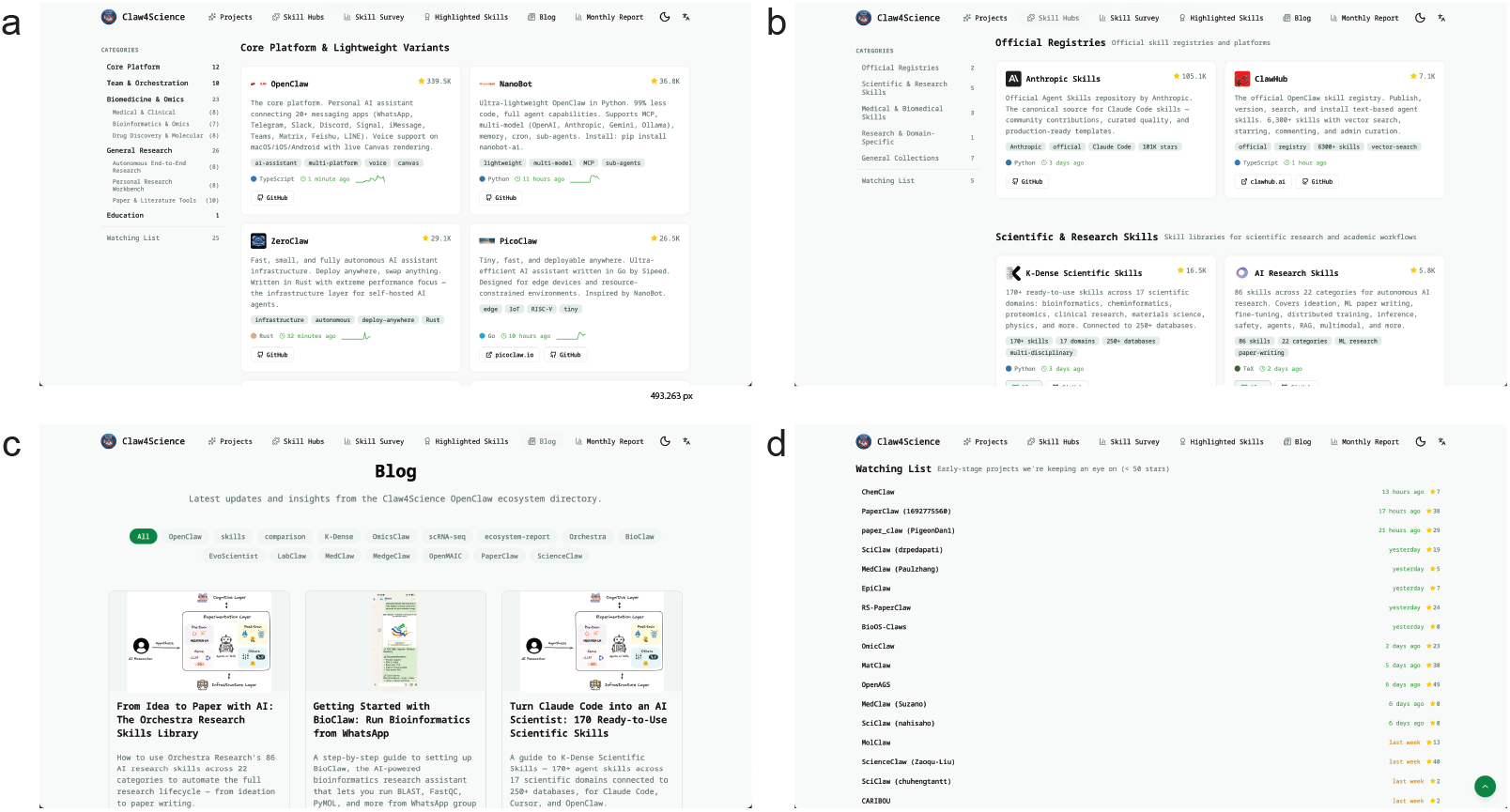
The Claw4Science platform. (a) Project directory view, which organizes OpenClaw-related projects into functional categories. (b) Skill hub interface, which aggregates offcial registries and community-maintained skill libraries into a unified navigation layer. (c) Blog and content view, which provides tutorials, updates, and ecosystem insights. (d) Watching list, which tracks emerging projects and recent activity across the ecosystem.

The dataset also shows that the ecosystem is not limited to domain-specific biology tasks. Many skills support general research activities such as literature review, academic writing, and data analysis. This suggests that the skill system operates across the full research workflow rather than only within specific domains. Overall, the skill layer shows both concentration and diversity. A few domains dominate in scale, while many smaller categories extend the system into new areas. This pattern is consistently observed in our dataset.

### 4.3. Why Skills Explain the Ecosystem’s Growth

Based on our dataset, the OpenClaw ecosystem shows a pattern of sustained community-driven expansion that is not observed in many other open-source agent frameworks. This pattern can be explained by several properties of the skill system.

The first is a low barrier to contribution. Writing a skill requires domain knowledge and the ability to describe a process clearly. It does not require programming or knowledge of a specific framework. Many more researchers can describe an analysis in natural language than implement it as a full software package. Similar patterns have been observed in low-code systems, where lowering technical requirements increases participation [47]. The second property is composability. Skills are modular and can be used together in a single session. A researcher can combine literature search, data retrieval, analysis, and visualization by calling different skills in sequence. This does not require additional integration work. The system manages the flow of information between steps.

The third property is transparency. Skills are written in plain text, so users can inspect what the agent will do before execution. This supports reproducibility in scientific computing [48]. Skills also act as both documentation and execution, which reduces missing details in workflows [41]. The fourth property is a positive feedback loop in the community. As more skills become available, the system becomes useful to more researchers. As more researchers use it, more of them contribute new skills. This cycle expands the coverage of the system over time. Similar dynamics have been described in platform ecosystems [51]. In this case, the process is fast because contributing a skill only requires writing a Markdown file and sharing it.

### 4.4. Formalizing the Skill Paradigm: The Claw4S Conference

As the skill ecosystem grows, evaluation becomes an increasingly important issue. Skills vary widely in quality, and there is no standard way to assess them. Based on our dataset, this variation is evident across repositories and domains. The Claw4S Conference, scheduled for 2026, is an early attempt to address this problem [3]. It proposes to review executable skill files instead of traditional papers, with criteria such as executability, reproducibility, scientific rigor, generalizability, and clarity. This reflects a broader shift toward treating skills as a unit of scientific contribution. This direction is still emerging. It remains unclear whether such evaluation systems can maintain quality while keeping participation open.

## 5. Challenges and Open Problems

The rapid growth of the OpenClaw ecosystem has introduced several challenges. These are not unique to this community, but they are more visible here due to the speed of growth and the scientific context. We highlight four main issues: naming collisions, skill quality, reproducibility, and the lack of benchmarks.

### 5.1. Naming Collisions and Ecosystem Governance

Naming collisions are common in the ecosystem. Based on our dataset, we identify 23 cases across 91 projects. The ScienceClaw name alone is used by four independent systems with different designs. Similar issues appear in MedClaw and SciClaw. As a result, users cannot easily identify which project they are using. To address this issue, the Claw4Science platform explicitly disambiguates projects with overlapping names by linking each entry to its corresponding repository and metadata. This allows users to compare similarly named systems directly and reduces confusion during navigation.

This problem reflects a broader lack of coordination. There is no naming convention, no registry, and no shared governance process. Governance gaps extend beyond naming. There is no standard for deprecating outdated projects or identifying misleading ones. The current solution relies on manual curation, which does not scale. More formal governance structures will be needed as the ecosystem grows.

### 5.2. Skill Quality Variance

Based on our dataset, skill quality varies widely across repositories and domains. Some skills are well-tested and reviewed, while others are incomplete or incorrect. The current classification system measures coverage, not quality. A skill can be correctly categorized but still produce unreliable results. This is a structural issue. The low barrier to contribution makes it easy to submit new skills, but there are few quality signals. Unlike traditional software, there are no standard mechanisms such as code review or usage metrics at scale. The problem is further amplified by hub fragmentation. Skills for the same task are often distributed across multiple repositories, with no clear guidance on which to use. This makes it diffcult for researchers to select reliable workflows.

### 5.3. Reproducibility Under External Dependencies

Based on our dataset, reproducibility is strongly affected by external dependencies. Skills that rely on APIs, models, or databases may produce different results over time. Model updates, API changes, and software version changes can all break a workflow or change its output. Containerization can help control the execution environment, but it does not resolve model drift. A system that calls external models will still change as those models evolve. Local-first approaches reduce this problem, but they require more resources and are not always practical. The core issue is that skills are static descriptions but depend on dynamic systems. This creates reproducibility risks that accumulate over time.

### 5.4. Benchmark Absence

There is no widely adopted benchmark for evaluating scientific skill-based systems in this ecosystem. Based on our dataset, this gap is particularly visible in bioinformatics and biomedical domains, where correctness, reproducibility, and tool integration are critical. Existing benchmarks evaluate general agent capabilities [1, 52], but they do not capture the properties that define scientific skill workflows. They are not designed for domain-specific settings such as bioinformatics or biomedical research.

A key direction for future work is to develop a benchmark tailored to scientific skills. Such a benchmark should evaluate portability, composability, and reproducibility across different environments. Our dataset provides a starting point for this effort, as it captures both the diversity of skills and their distribution across scientific domains. Without such benchmarks, it is diffcult to assess which systems are reliable for scientific use.

## 6. Discussion and Conclusion

This work contributes three components to the OpenClaw scientific ecosystem: a curated dataset, a systematic analysis, and a platform for exploration and use. We construct a dataset of OpenClaw-related projects and skills, and use it to analyze the structure of the ecosystem, including its layered architecture, application domains, and cross-cutting patterns. Our analysis shows that the growth of the ecosystem is driven by a simple but powerful design choice. The skill system lowers the barrier to contribution and allows workflows to be shared across different runtimes. This leads to rapid expansion and broad coverage of scientific tasks.

At the same time, the ecosystem is still early in its development. Many projects are small and specialized. Based on our dataset, quality varies across skills, and reproducibility is not guaranteed when external dependencies are involved. Naming inconsistencies and the lack of governance make navigation diffcult. The absence of benchmarks also limits systematic evaluation. These issues suggest that the ecosystem has not yet reached a stable or standardized state.

To address these challenges, we develop Claw4Science (https://claw4science.org), a platform built on top of our dataset. It organizes projects and aggregates scientific skill hubs into a unified interface (Figure 4). The platform provides disambiguation for overlapping project names and improves navigation across distributed repositories. It is designed for scientific use, with a focus on bioinformatics and biomedical workflows, and serves as a practical entry point for exploring the ecosystem.

Looking forward, several directions are important for the next stage of this ecosystem. First, evaluation frameworks are needed to compare skills and agent systems in a consistent way. Our dataset provides a starting point for developing such benchmarks. Second, governance structures will be necessary to manage naming, quality, and project lifecycle. Third, reproducibility will require better control of model and data dependencies. Finally, lightweight and edge-deployed agents remain underexplored, despite their potential for real-world scientific use.

In summary, the OpenClaw ecosystem represents an early but important step toward a more modular and shareable model of scientific computation. This work provides a dataset and platform that make this ecosystem more accessible and analyzable, especially for scientific workflows. Its future impact will depend on whether the community can balance openness with reliability.

## Supplementary Materials

### A. Detailed Methodology

#### A.1. Project Discovery

We collected candidate projects through GitHub keyword search, social media monitoring, community recommendations, and cross-references. Each project was manually reviewed to ensure relevance to scientific workflows and public availability.

#### A.2. Skill Survey Pipeline

We developed an automated pipeline to scan 12 science-focused repositories via the GitHub API. The pipeline recursively identified skill files and classified them into 34 categories using keyword-based matching.

#### A.3. Name Disambiguation

We manually identified naming collisions across projects and grouped them into distinct clusters.

#### A.4. Data Maintenance

The dataset is updated periodically. The latest version is available at https://claw4science.org.

### B. List of Curated OpenClaw Projects

**Table 1.**
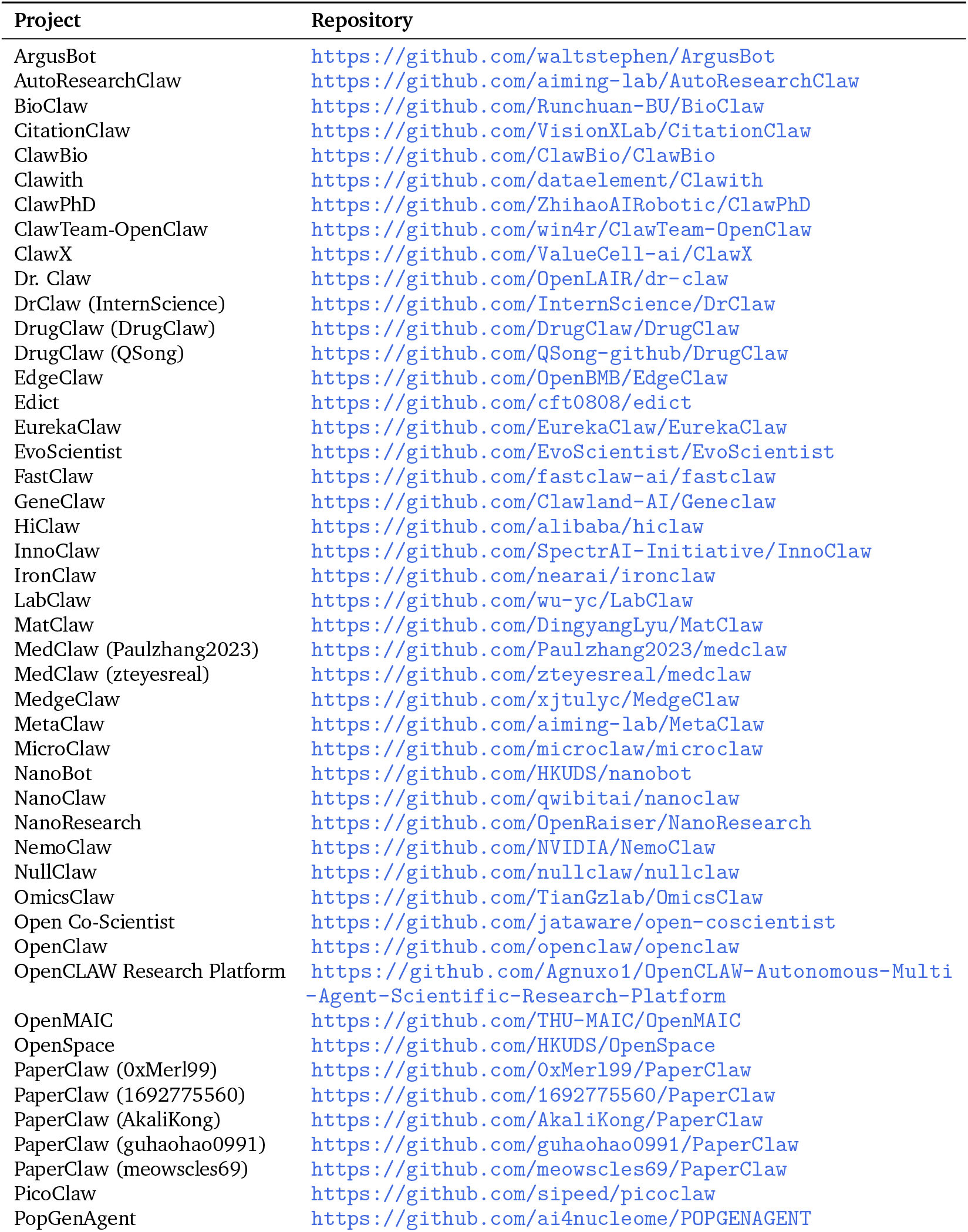
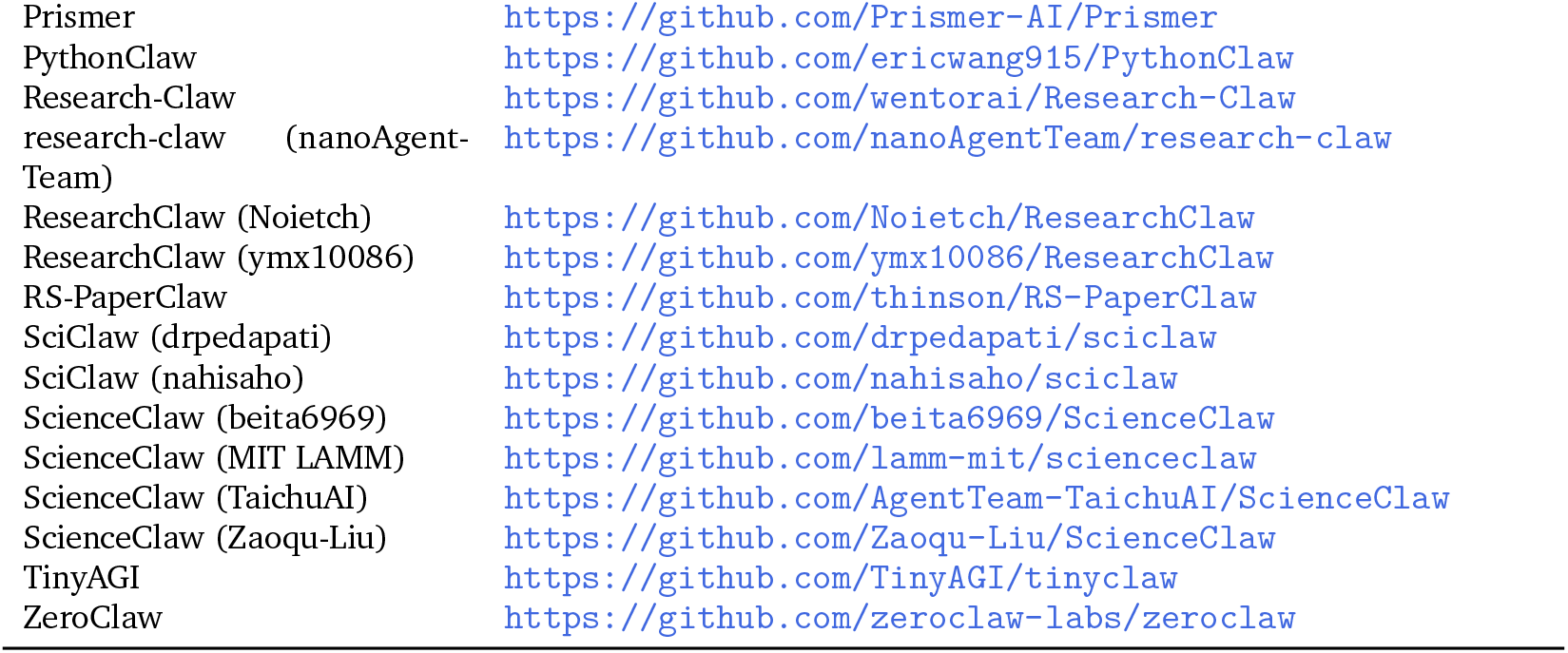
Curated list of projects included in the post-OpenClaw ecosystem analysis.

### C. Case Study: BioClaw in the OpenClaw Ecosystem

To demonstrate the practical utility of the Claw4Science platform and its connection to the OpenClaw ecosystem, we present a case study using BioClaw, a domain-specific agent in the bioinformatics branch. As shown in Figure 2, BioClaw is positioned within the bioinformatics application layer of the post-OpenClaw ecosystem, built on top of OpenClaw-compatible runtimes (e.g., NanoClaw and related lightweight variants). In this setting, BioClaw does not operate as a standalone system, but as a composition of reusable skills discovered through the Claw4Science platform.

We evaluate this setup on a realistic biological task: protein sequence analysis from raw amino acid input. Given a protein sequence, BioClaw dynamically retrieves and executes relevant skills, including sequence analysis, homology search, structure mapping, and functional annotation. The system first performs a BLASTP search against Swiss-Prot and identifies the sequence as human *γ*-tubulin (TUBG1), with near-complete coverage and essentially perfect identity. Sequence alignment confirms that the core residues match the canonical TUBG1 protein, with additional C-terminal residues corresponding to a TEV cleavage site and His-tag from recombinant expression.

BioClaw then maps the sequence to the AlphaFold structural model and highlights the experimentally annotated GTP-binding region (residues 142–148). As shown in Figure 5, the predicted structure exhibits high confidence across most residues, consistent with the known tubulin fold. Functionally, *γ*-tubulin is a core component of the *γ*-tubulin ring complex (gTuRC) and plays a critical role in microtubule nucleation. The correct identification of this protein, along with recovery of its key functional region, demonstrates that BioClaw can produce biologically meaningful interpretations.

**Figure 5.**
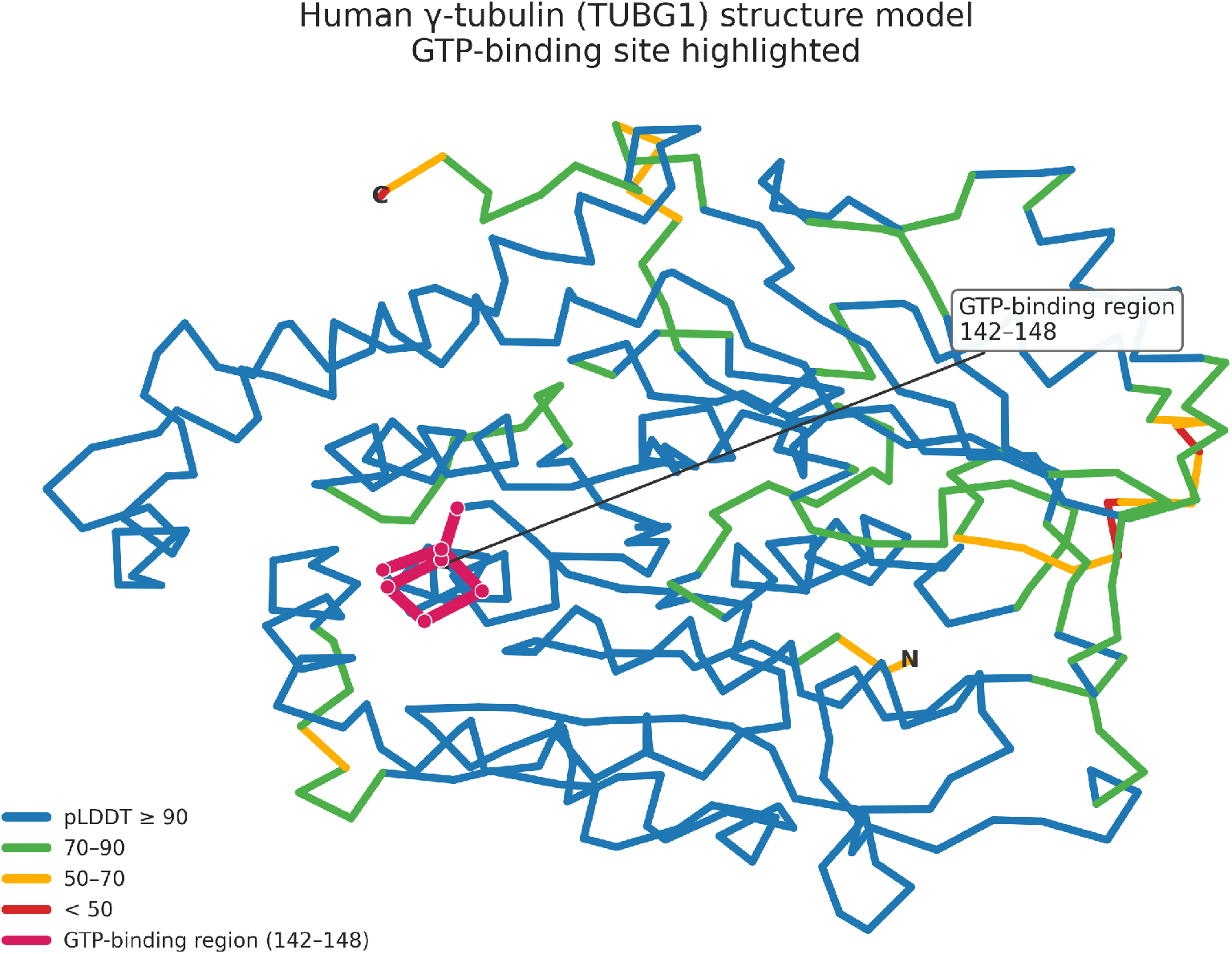
Example of a BioClaw-enabled analysis within the OpenClaw ecosystem. Given an input protein sequence, the system integrates distributed skills for sequence analysis, structural prediction, and functional annotation. The predicted structure corresponds to human *γ*-tubulin (TUBG1), with the backbone colored by AlphaFold confidence (pLDDT) and the experimentally annotated GTP-binding region (residues 142–148) highlighted in magenta. This example illustrates how ecosystem-level composition of reusable skills enables end-to-end scientific workflows.

In addition to sequence-to-structure analysis, we further evaluate BioClaw on data-driven transcriptomics tasks. Specifically, we consider a publicly available RNA-seq dataset (GSE150316) and perform differential expression analysis. BioClaw retrieves the dataset, applies filtering and normalization, conducts statistical testing with Benjamini–Hochberg FDR correction, and generates a publication-quality volcano plot (Figure 6). The analysis identifies significantly upregulated and downregulated genes, with strong signals corresponding to viral transcripts and host response patterns. This demonstrates that BioClaw can extend beyond sequence-level analysis to support end-to-end data-driven biological workflows.

**Figure 6.**
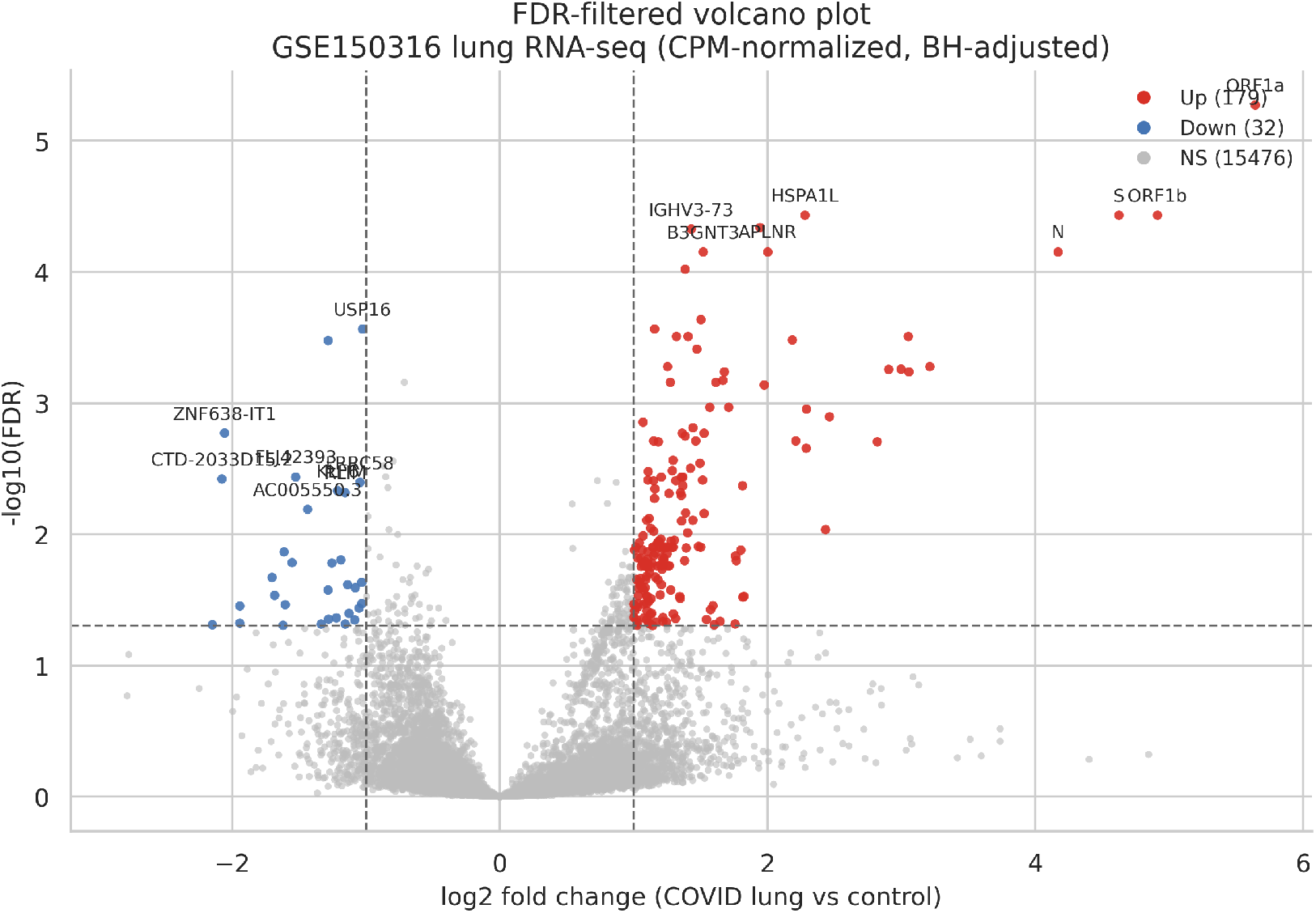
BioClaw-enabled RNA-seq differential expression analysis on a publicly available dataset (GSE150316). The system retrieves the dataset, performs differential expression analysis with FDR correction, and generates a publication-quality volcano plot. Significantly upregulated genes are shown in red and downregulated genes in blue (|log_2_ FC| *>* 1, FDR *<* 0.05). Representative genes are labeled. This example demonstrates how BioClaw integrates data retrieval, statistical analysis, visualization, and biological interpretation within the OpenClaw ecosystem.

Importantly, these results are not produced by a monolithic system, but emerge from the composition of distributed skills within the OpenClaw ecosystem, discovered and organized through the Claw4Science platform. This illustrates how ecosystem-level design enables real scientific workflows by combining reusable, decentralized components. Overall, this case study shows that Claw4Science serves not only as a dataset and directory, but also as an enabling layer for executing end-to-end scientific analysis through ecosystem-integrated agents such as BioClaw. Similar behavior is observed across additional tasks (not shown).

## Notes

### Competing Interest Statement

The authors have declared no competing interest.

